# Epigenetic modulation of selected immune response genes and altered functions of T lymphocytes and macrophages collectively contribute to autoimmune diabetes protection

**DOI:** 10.1101/2021.07.15.452572

**Authors:** Sundararajan Jayaraman, Maria Arianas, Arathi Jayaraman

## Abstract

We have previously demonstrated that treatment of female NOD mice with the histone deacetylase inhibitor Trichostatin A (TSA) bestowed irreversible protection against autoimmune diabetes. Herein we show that drug treatment diminished the infiltration of the pancreas with CD4^+^ and CD8^+^ T cells and Ly-6C^+^ monocytes. Significantly, TSA administration selectively repressed the expression of a set of genes exaggerated during diabetes and constitutively expressed primarily in the spleen and rarely in the pancreas. These genes encode lymphokines, macrophage-associated determinants, and transcription factors. Although the copy numbers of many histone deacetylases increased during diabetes in the spleen and pancreas, only those upregulated in the spleen were rendered sensitive to repression by TSA treatment. The T lymphocytes derived from drug-treated donors displayed diminished diabetogenic potential following transfer into immunodeficient NOD.*scid* mice. In the immunocompromised recipients, diabetes caused by the transfer of activated T lymphocytes from untreated diabetic mice was hampered by the co-transfer of highly purified splenic Ly-6C^+^ macrophages from drug-treated mice. However, the transfer of Ly-6C^+^ macrophages from drug-treated mice failed to block ongoing diabetes in wild-type NOD mice. These data demonstrate that the modified gene expression and functional alteration of T lymphocytes and macrophages collectively contribute to diabetes protection afforded by the histone modifier in female NOD mice.

## 1. Introduction

Type 1 diabetes (T1D), a T cell-mediated autoimmune disease, occurs at an alarming rate in genetically susceptible individuals [1–3]. Although genome-wide association studies implicated more than 40 different genetic loci, the causal genes of T1D remain unknown [4]. Lack of familial history and robust concordance among identical twins in developing T1D implied the involvement of environmental factors. Epigenetics provides a mechanism by which external factors can produce a variety of phenotypic variations with identical genotypes. Epigenetic mechanisms include DNA methylation, post-translational modifications of histones, and microRNA-mediated gene regulation [5]. Methylation of cytosine residues in gene promoters results in transcriptional silencing of some [6] but not all genes [7]. Reversible acetylation of the e-amino group of lysine in the histone tails by histone acetyltransferases and deacetylation by histone deacetylases (HDACs) are the best-characterized post-translational modifications of histones [8]. Transcriptional permissiveness correlates with histone acetylation in contrast to deacetylation resulting in gene repression. Non-selective, small molecule HDAC inhibitors can induce chromatin modifications and thereby affect gene expression. Trichostatin A (TSA) derived from *Streptomyces platensis* is the most potent inhibitor of class I, class IIa, class IIb, and class IV HDACs, which increases histone acetylation resulting in up- and down- regulation of genes *in vitro* [9] and *in vivo* [10–16].

Previously, we have demonstrated the utility of TSA to mitigate autoimmune diseases such as T1D [10–13] and experimental autoimmune encephalomyelitis [14–16]. The protection against T1D afforded by the histone modifier was accompanied by reduced inflammation of the pancreatic islets and regeneration of insulin-producing β-cells [10]. Significantly, treatment with TSA abrogated the ability of splenocytes to adoptively transfer T1D into immunodeficient NOD.*scid* mice, indicating the effect of the epigenetic drug on diabetogenic T lymphocytes [11]. Transcriptome analysis of *uninduced* splenocytes unraveled the exaggerated expression of inflammatory genes, including the macrophage selective *Cela3b* (elastase 3) in diabetic mice repressed by TSA. These results are consistent with the possibility that chromatin remodeling by TSA can retard the induction and progression of T1D by altering the transcriptional programs of multiple cell types. In the current investigation, we analyzed the effects of the epigenetic drug on T lymphocytes and macrophages, respectively implicated in induction and manifestation of T1D [17–19], by quantitative reverse transcriptase-mediated polymerase chain reaction (qRT-PCR) and an adoptive transfer model. Our study unraveled that a select set of both exaggerated and constitutively expressed genes encoding cytokines, cell surface determinants, transcription factors, and HDACs was modulated by the histone modifier more predominantly in the spleen. Our data also demonstrate the distinct effects of TSA on the function of T-cells and macrophages. These data lend novel insights into the mechanisms of T1D and its possible manipulation by epigenetic reprogramming.

## 2. Materials and materials

### 2.1. Diabetes treatment

The Office of Animal Care and Institutional Biosafety of the University of Illinois at Chicago approved the animal protocol. Experiments were conducted following the NIH guide for the care and use of laboratory animals.

Six to eight weeks old female NOD/ShiLtj (H-2^g7^) mice purchased from The Jackson Laboratory (Bar Harbor, ME) were maintained under specific pathogen-free conditions and provided with animal chow and water *ad libitum*. Mice were injected s.c. with 500 μg/Kg body weight of TSA or DMSO at weekly intervals between 16 and 24 wks of age [10–11]. Non-fasting blood glucose level exceeding 250 mg/dL on two consecutive weekly determinations was considered as diabetic.

### 2.2 Confocal microscopy

The following antibodies were used for staining pancreatic sections: guinea pig anti-insulin Ab (Zymed Laboratories, South San Francisco, CA), tetramethylrhodamine iso-thiocyanate–rabbit antisera raised against guinea pig Ig (Sigma Aldrich, St. Louis, MO), anti-Ly-6C-FITC (clone HK1.4, eBiosciences, San Diego, CA), anti-Ly-6G-FITC (clone 1AB, BD Pharmingen, San Diego, CA), rat monoclonal anti-CD4 (clone RM4-5, eBiosciences-ThermoFisher Scientific, Waltham, MA), DyLight™ 488 goat anti-rat IgG (catalog # 405409, BioLegend, San Diego, CA) and anti-CD8-FITC (clone 53-6.7, eBiosciences). Confocal images were acquired using a Zeiss LSM510 laser scanning microscope and processed using the Zeiss LSM Image browser (4.0 version; Zeiss, Oberkochen, Germany) and Adobe Photoshop Elements version 9.0, as described [20].

### 2.3 Gene expression analysis

Spleens were harvested from 20-24-wks old diabetic mice and 26-28 wks old mice treated with TSA from 16 to 24-wks of age. Total RNA was extracted from the spleen and pancreas of five *individual mice* per group/experiment, treated with Turbo DNase, and converted to cDNA using High-capacity cDNA Reverse Transcription kit (Applied Biosystems), as described [10–11, 14–16, 20]. Real-time qRT-PCR was performed in *triplicate* on Applied Biosystems ViiA7 Real-time PCR system using 1 μl of cDNA equivalent to 100 ng of total RNA and 2X SYBR Green master mix. The primers used in this study were designed and validated previously [10–11, 14–16, 20] and synthesized at the Integrated DNA Technologies (Coralville, IA). The level of specific gene expression in each sample was ascertained using *Gapdh* as the normalizer and the 2^-ΔΔ*C*^T method.

### 2.4 Adoptive transfer of diabetes

Spleens were harvested from 20-24-wks old diabetic mice and from 26-28 wks old mice treated with TSA between 16 to 24-wks of age. Splenocytes (10 × 10^6^/ml) were cultured with five μg of Concanavalin A (Sigma Aldrich) for two days as described [19]. More than 95% of these cells were CD3^+^, as determined by flow cytometry. Splenocytes were also stained with PE-Cyanine 5 conjugated anti-CD11b (clone M1/70, eBiosciences) and anti-Ly-6C-FITC antibodies and sorted on a MoFlo Astrios cell sorter, which yielded 2.27 % of the input with > 98% CD11b^+^Ly-6C^+^ cells. Both male and female NOD.*scid* mice were injected i.v. with activated 2 × 10^6^ T lymphocytes with or without sorted 2 × 10^5^ CD11b^+^Ly-6C^+^ cells or purified monocytes alone. Splenocytes were also stained with anti-Ly-6C-FITC and incubated with anti-FITC antibody and purified on a magnetic column (Miltenyi Biotec Inc., Auburn, CA). This procedure yielded 2 % of the input with >95% Ly-6C^+^ cells, as determined by flow cytometry. Female wild-type NOD mice received two injections of purified 2 × 10^5^ Ly-6C^+^ cells at 13 and 21-wk of age. Numbers of mice tested are given in the figure legends.

### 2.5 Statistics

The difference in diabetes incidence between controls (untreated/DMSO-treated mice) and those treated with TSA was analyzed for statistical significance using the Wilcoxon Signed Rank Test. Gene expression data were analyzed for statistical significance between indicated groups by two-way ANOVA or two-tailed unpaired *t*-test using GraphPad Prism (6.0) software and indicated in individual figure legends. A *P*-value of <0.05 was considered significant.

## 3. Results

### 3.1 Protection from T1D is associated with constrained infiltration of inflammatory cells into the islets

We have consistently observed that 80-100% of female NOD mice procured from The Jackson Laboratory and housed in our animal facility developed T1D by 18 weeks of age [10–13, 20]. Weekly treatment with TSA starting at 16 wk and until 24 wks of age substantially attenuated T1D (Supplementary Fig. 1). Injection of the vehicle DMSO failed to alter the course or intensity of the disease [10–13]. We have also documented that DMSO treatment did not change gene expression compared to untreated diabetic mice. Since DMSO-treated mice did not have noticeable changes in disease course and gene expression compared to untreated diabetic mice, we compared the litter mates of sex- and age-matched untreated diabetic mice with TSA-treated mice in all experiments. Treatment with TSA reduced the invasive cellular infiltration of the islets and reversed moderate to a severe loss of insulin-producing beta cells [10]. However, the nature of cellular infiltration was not previously characterized. To this end, we performed confocal analysis, which indicated the distribution of Ly-6C^+^ macrophages, Ly-6G^+^ neutrophils, CD4^+^, and CD8^+^ T-cells across the entire area of the diabetic pancreas (Fig. 1). Treatment with TSA significantly reduced the infiltration of primarily CD4^+^ cells and, to some extent, CD8^+^ T-cells implicated in T1D [17]. The influx of Ly-6C^+^ macrophages but not neutrophils was also reduced by TSA treatment, consistent with a role for macrophages in diabetes [18–19]. Flow cytometric analysis confirmed that the numbers of CD11b^+^Ly-6C^+^ monocytes but not CD11b^+^F4/80^+^ macrophages or CD11c^+^ dendritic cells were also diminished in the spleen of TSA-treated mice (Supplementary Fig. 2). These data indicated that reduced influx of CD4^+^ and CD8^+^ T cell subsets and Ly-6C^+^ macrophages into the islets serves as a mechanism of TSA-mediated amelioration of autoimmune diabetes.

**Fig. 1.**
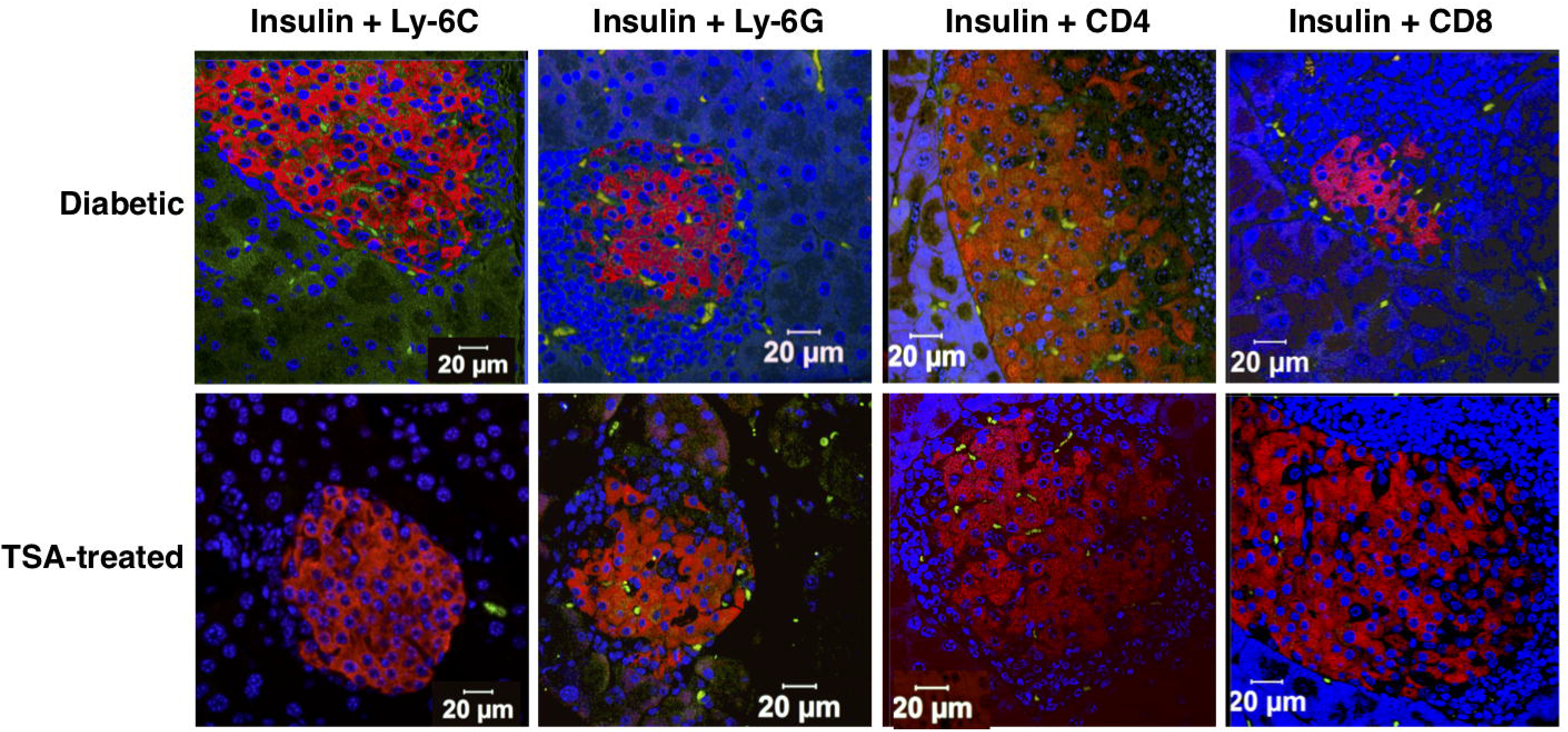
Cellular infiltration in the pancreas was retarded by TSA treatment. Diabetic mice and those treated with TSA during 16 and 24-wk of age were sacrificed when they were 26-28-wks old. Sections of the pancreas were stained with guinea pig antisera against insulin followed by TRITC labeled rabbit anti-guinea pig antibody (red), along with FITC-conjugated anti-Ly-6C, anti-Ly-6G, anti-CD4 or anti-CD8 antibodies (green). After counterstaining with Hoechst (blue), the sections were analyzed on a confocal microscope. Representative images from multiple experiments are shown.

### 3.2 TSA treatment selectively repressed the expression of Th1 and Th17 lymphokines, primarily in the spleen

Our previous transcriptome analysis of *uninduced* splenocytes indicated the downregulation of a few inflammatory genes, including *Cela3b* (elastase) by TSA treatment [11]. In the current investigation, we analyzed a total of 43 genes selected for their putative roles in T1D. These included genes encoding 11 lymphokines, 13 accessory cell-associated determinants, seven transcription factors, a chemokine, and 11 genes encoding HDACs. Total RNA was extracted from diabetic and TSA-treated animals, converted to cDNA, and used as the template in the qRT-PCR assay as described before [10–16]. The genes critical for Th17 cell development, namely *Il17a* and *Il23* [21–22], and the Th1-specific *Ifng* [23] were significantly increased in the spleen of diabetic mice compared to the prediabetic mice displaying normal levels of blood glucose (Fig. 2a). Significantly, TSA treatment diminished the levels of these genes. Although *Il4*, *Il18*, and *Il27p28* were not exaggerated during diabetes, they were nevertheless repressed by treatment with the histone modifier. However, other genes such as *Il22*, *Il27ebi3*, *Il10*, *Il12*, and *Tgfb* were neither upregulated during diabetes nor modulated by the histone modifier, indicating that they are unlikely to contribute to diabetogenesis in this model.

**Fig. 2.**
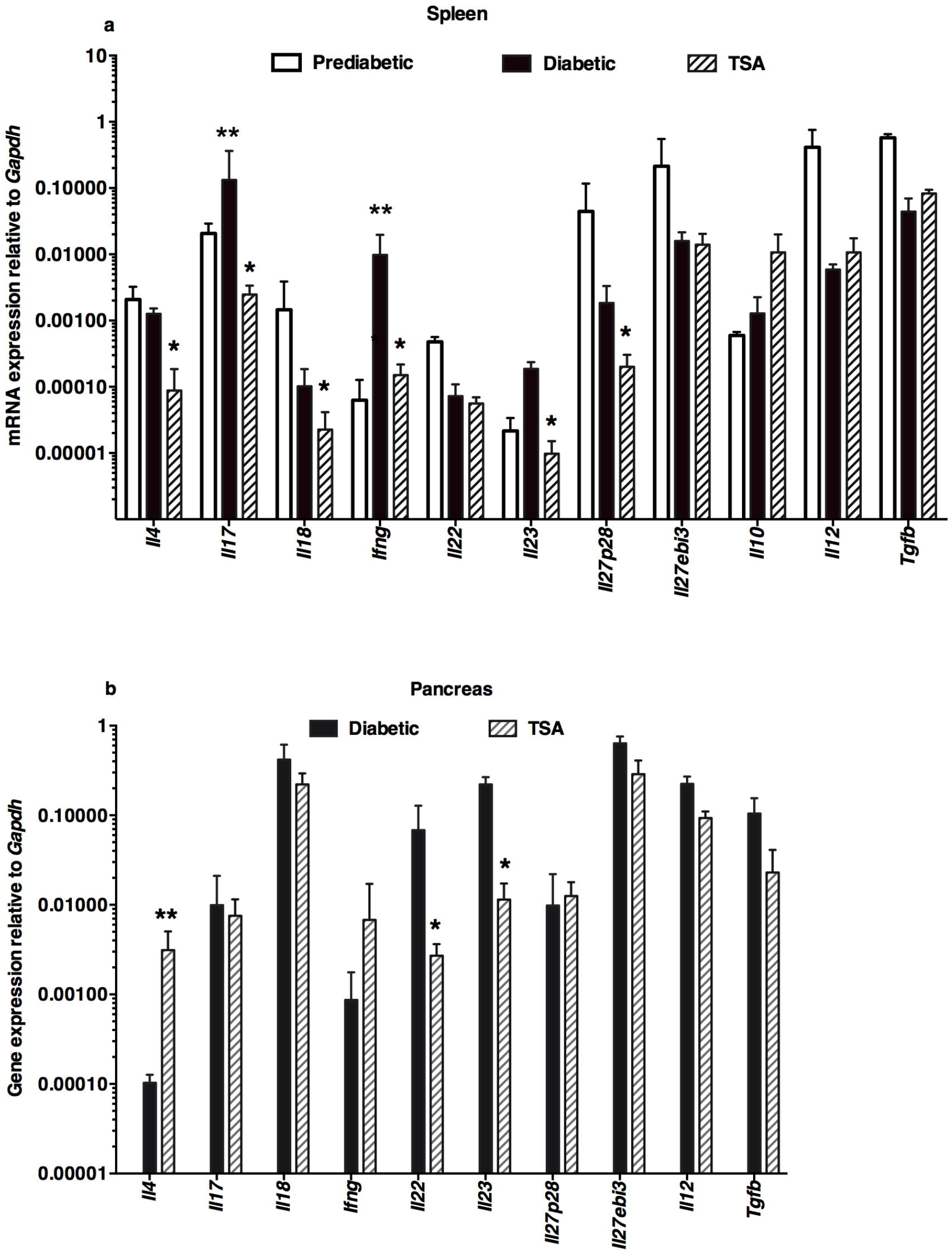
Differential modulation of lymphokine genes by the histone modifier. Spleen was harvested from 12-wk old prediabetic mice and 25-26-wks old diabetic mice. Pancreas was collected from 25-26-wks old diabetic mice. Mice treated with TSA between 16 and 24-wks of age were sacrificed when they were 25-26-wks old and spleen and pancreas were harvested. Five mice per group were tested in three different experiments. Total RNA was extracted from the spleen (**a**) and pancreas (**b**) of five individual mice was analyzed, converted to cDNA, and used as the template in qRT-PCR. Data indicate the mean +/− SD. Representative data from three independent experiments are depicted. Asterisks indicate statistical significance in the spleen samples from prediabetic, diabetic, and TSA-treated mice, as assessed by two-way ANOVA (**a**) and by two-tailed unpaired *t*-test between diabetic and TSA-treated pancreas (**b**) (*P*<0.05). Increased values between prediabetic and diabetic mice were indicated by double asterisks whereas a single asterisk denoted the decrease in values in TSA-treated mice compared to diabetic mice.

Preliminary data indicated that the expression of the genes analyzed herein remained comparable in the pancreas of prediabetic and overtly diabetic mice. Therefore, the gene expression levels in the pancreas of diabetic and TSA-treated mice were compared. Unlike the spleen, TSA treatment increased the level of *Il4* but repressed the transcription of *Il22* and *Il23* in the pancreas (Fig. 2b). Collectively, these results suggest that TSA treatment differentially regulated the expression of both inducible and constitutively expressed immune response-related genes in the spleen and pancreas, and inversely correlated with normoglycemia.

### 3.3 Histone modifier regulates constitutive expression of macrophage-associated cytokines primarily in the spleen

The chemokine CCL2, implicated in the mobilization of myeloid cells from the bone marrow causing insulitis [24], was upregulated in the spleen of diabetic mice, which was repressed by TSA treatment (Fig. 3a). Although macrophage-selective genes such as *Csf2*, *Mmp12*, and *Ym1* were not exaggerated during diabetes and yet were reduced by the histone modifier. Significantly, *Cela3b* expression was lower in TSA-treated spleens, as shown earlier [11]. Only the steady-state level of *Cd39* was subject to regulation by the HDAC inhibitor in the pancreas (Fig. 3b). These data suggest that TSA treatment predominantly diminished the constitutive expression of many macrophage-selective genes in the spleen compared to the pancreas.

**Fig. 3.**
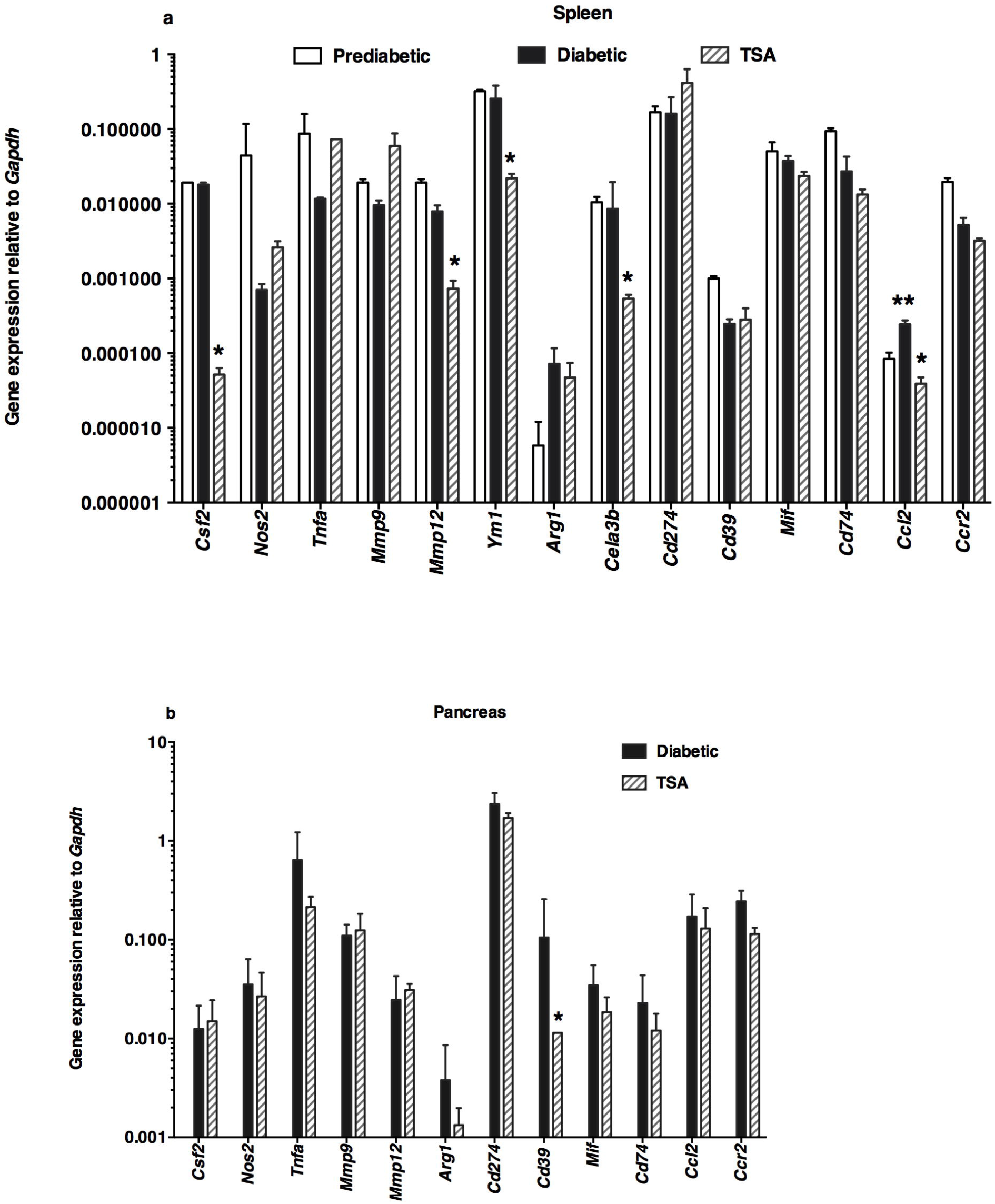
Differential regulation of accessory cell-associated determinants following TSA treatment. The expression levels of genes associated with accessory cells were analyzed in (**a**) the spleen and (**b**) pancreas. Total RNA was extracted and gene expression was analyzed by qRT-PCR using the same cDNA preparations analyzed in Fig. 2. Data indicate arithmetic mean +/− SD of three determinations. Representative data from three independent experiments are depicted (*n*=5 mice per group/experiment). Asterisks indicate statistical significance in the spleen samples from prediabetic, diabetic, and TSA-treated mice, as assessed by two-way ANOVA (**a**) and by two-tailed unpaired *t*-test between diabetic and TSA-treated pancreas (**b**) (*P*<0.05). Increased values between prediabetic and diabetic mice were indicated by double asterisks, whereas the decrease in values in TSA-treated mice compared to diabetic mice was denoted by a single asterisk.

### 3.4 Epigenetic drug impacts the expression of transcription factors sparingly in the spleen

Notably, the transcription factor *Ahr* involved in nitric oxide and arginine production, and alteration of M1/M2 macrophage polarization [25] was transcriptionally increased in the spleen of diabetic mice and reduced following TSA treatment (Fig. 4a). Only the constitutive expression of *Rorgt*, crucial for the transcription of IL-17A [26], was repressed by TSA treatment. Surprisingly, *Tbet* and *Gata3*, respectively involved in the transcription of IFN-γ and IL-4, and other transcription factors such as *Eomes*, *Dec1*, and *Foxp3* examined were neither transcriptionally upregulated during diabetogenesis nor subject to epigenetic regulation. Interestingly, the *Foxp3* transcript level was lower in the TSA-treated pancreas (Fig. 4b). These data suggest that TSA-mediated protection from diabetes is associated with selective repression of IL-17A transcription in the lymphoid organ.

**Fig. 4.**
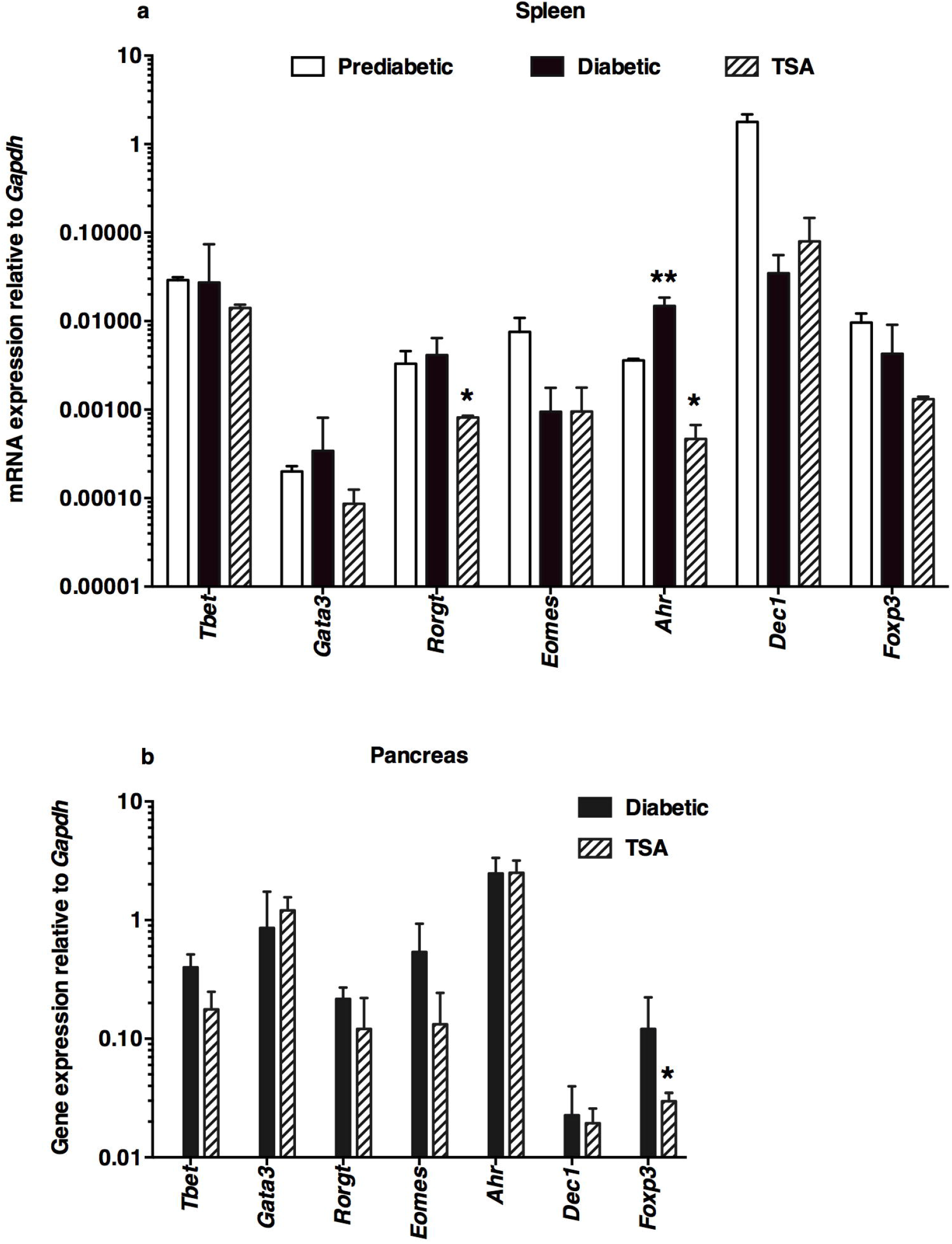
Selective regulation of the transcription factor genes by TSA. The expression levels of genes encoding various transcription factors were analyzed in (**a**) the spleen and (**b**) pancreas by qRT-PCR using the same cDNA preparations analyzed in Fig. 2 and 3. Data indicate the mean +/− SD of three determinations. Representative data from three independent experiments are depicted (*n*=5 mice per group/experiment). Double asterisks indicate a statistically significant increase between prediabetic and diabetic mice (\**P*<0.05), whereas a single asterisk denotes a decrease in the value between diabetic and TSA-treated groups (\*\**P*<0.01). Statistical significance was assessed respectively by two-way ANOVA (**a**) and two-tailed unpaired *t*-test between diabetic and TSA-treated pancreas (**b**) as directed by the GraphPad Prism software.

### 3.5 Epigenetic modifier differentially modulated HDACs in the spleen and pancreas

Although *in vivo* treatment of NOD mice with TSA induced the hyperacetylation of histone H3 in the spleen and pancreas of NOD mice [11], it is unclear whether it can differentially affect the expression of class I (*Hdac1*, *Hdac2*, *Hdac3*, and *Hdac8*), class IIa (*Hdac4*, *Hdac5*, *Hdac7*, and *Hdac9*), class IIb (*Hdac6* and *Hdac10*) or class IV (*Hdac11*) HDAC genes in the lymphoid system and the target organ pancreas. In the spleen of prediabetic, normoglycemic, 12-wk old female NOD mice, the basal levels of the *Hdac5* and *Hdac8* were minimal (Fig. 5a). During diabetes manifestation, the transcription of *Hdac4*, *Hdac8*, and *Hdac9* genes was increased, which was repressed by TSA treatment. Interestingly, *Hdac1*, *Hdac3*, *Hdac4*, *Hdac8*, and *Hdac11* were upregulated in the pancreas of diabetic mice (Fig. 5b). Whereas most of these genes were not repressed by the histone modifier, *Hdac4* was upregulated in drug-treated mice. It is noteworthy that *Hdac2*, *Hdac5*, *Hdac7*, *Hdac9*, and *Hdac10* genes could not be reliably detected in the pancreas of prediabetic, overtly diabetic, or TSA-treated mice by qRT-PCR in repeated experiments (n=25-30 mice). Thus, TSA-mediated protection against T1D was accompanied by reduction of the exaggerated expression of *Hdac4*, *Hdac8*, and *Hdac9* genes in the spleen, while *Hdac4* was upregulated in the pancreas.

**Fig. 5.**
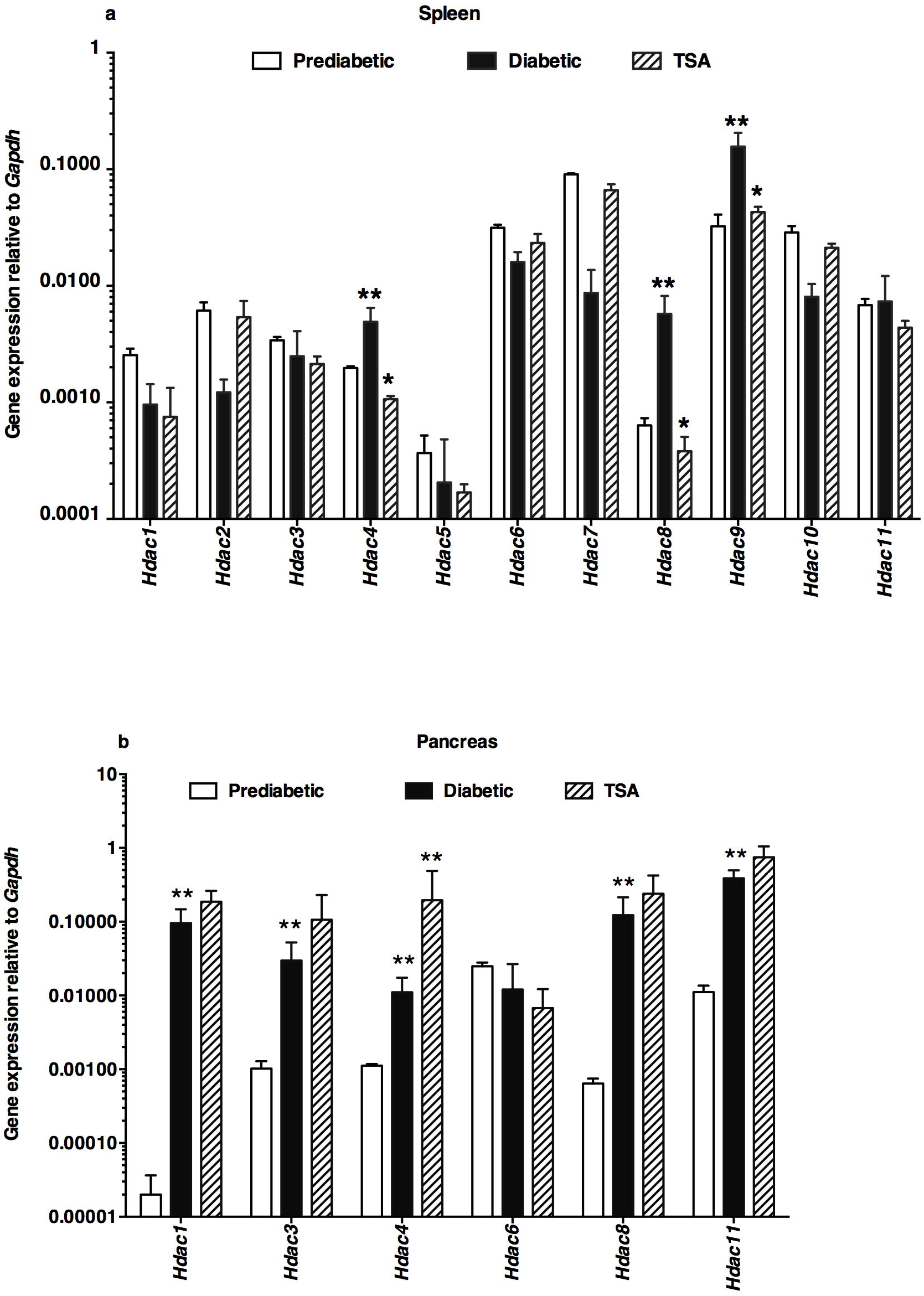
Differential modulation of *Hdac* genes by the epigenetic drug. The expression levels of *Hdac* genes in the spleen (**a**) and pancreas (**b**) were determined in cDNA samples illustrated in Fig. 2-4. Data indicate the mean +/− SD of three determinations. Representative data from three independent experiments are depicted (*n*=5 mice per group/experiment). An increase in the values between prediabetic and diabetic mice was indicated by double asterisks. The decrease in values in TSA-treated mice compared to diabetic mice was denoted by a single asterisk. Statistical significance was assessed using two-way ANOVA (**a**) and by a two-tailed unpaired *t*-test between diabetic and TSA-treated pancreas (**b**). \**P*<0.05 and \*\**P*<0.01.

### 3.6 TSA treatment differentially affected the function of T lymphocytes and macrophages

We next determined whether epigenetically modulated genes could impact the diabetogenic potential of T lymphocytes and the presumptive effector functions of macrophages in diabetes [18–19]. The data shown in Fig. 6a indicate that the transfer of Concanavalin A-activated T lymphocytes from diabetic mice mediated the disease in 100% of the immunodeficient NOD.*scid* mice by nine wks. However, similarly activated T lymphocytes derived from TSA-treated mice failed to induce the disease in NOD.*scid* recipients, consistent with the data obtained using whole splenocytes [11]. Interestingly, the adoptive transfer of sorted CD11b^+^Ly-6C^+^ macrophages from diabetic or TSA-treated mice did not cause diabetes in the immunodeficient mice, indicating that the macrophages are not the effectors of T1D in contrast to a previous observation indicating the beta-cell cytotoxicity *in vitro* by macrophages derived from diabetic mice [19]. Surprisingly, the co-transfer of CD11b^+^Ly-6C^+^ macrophages purified from the TSA-treated but not diabetic mice suppressed the T-cell mediated diabetes in NOD.*scid* recipients. In contrast to the data obtained in an adoptive transfer model, two consecutive injections of splenic Ly6-C^+^ macrophages purified from TSA-treated or diabetic mice failed to impede the ongoing disease in wild-type female NOD mice (Fig. 6b), inconsistent with diabetes suppression mediated by polarized M2-type macrophages reported previously [27]. These data suggest that the epigenetic modulation of macrophages resulted in distinct outcomes dictated by the recipients’ immune status.

**Fig. 6.**
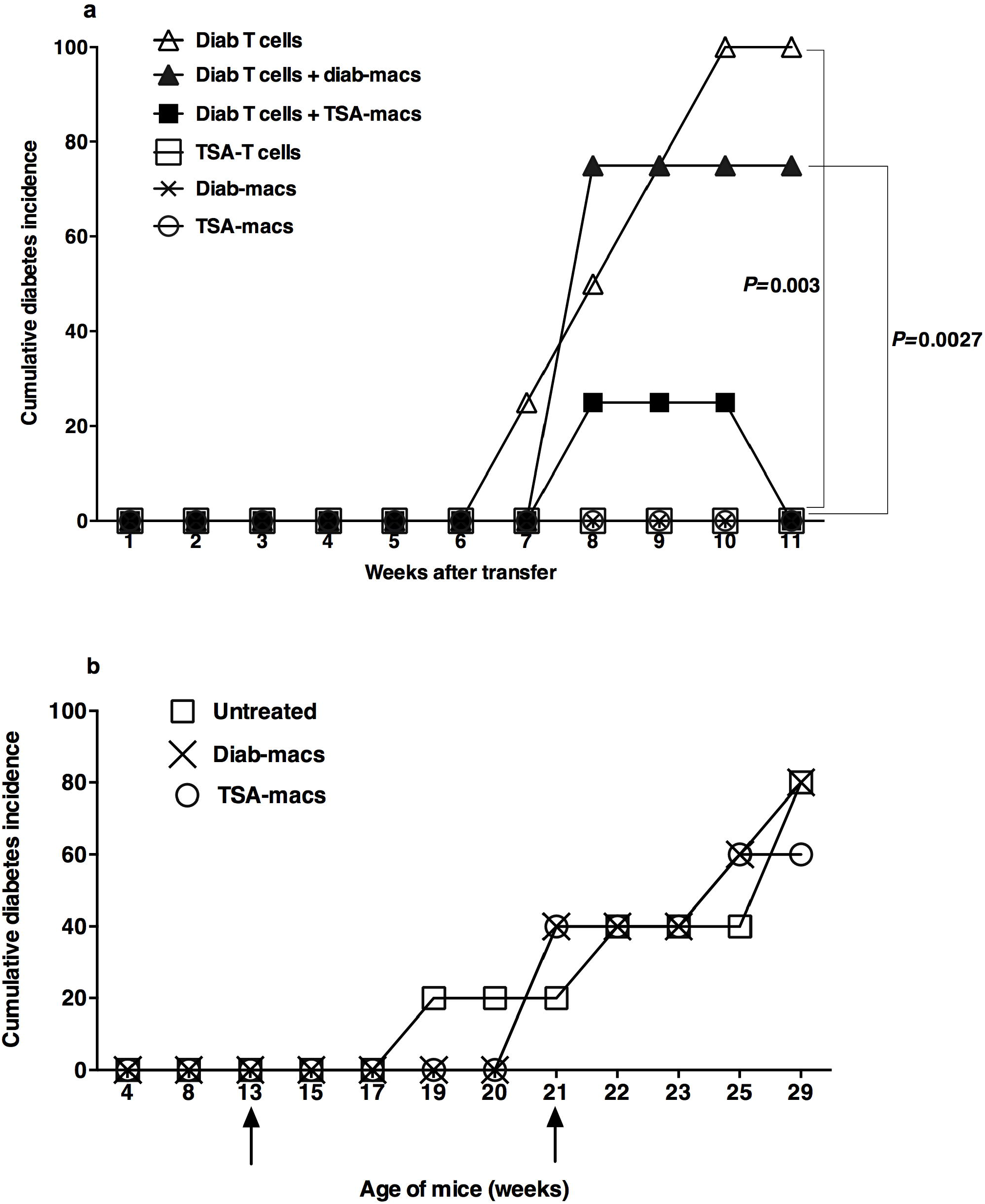
Epigenetic regulation of T cells and macrophages. (**a**) Diabetes was monitored in NOD.*scid* mice (10-12-wks old) injected i.v. with Concanavalin A-activated 2 × 10^6^ T lymphocytes from diabetic or TSA-treated NOD mice (26-28-wks of age) (*P*=0.0027). Mice were also co-transferred with diabetic T-cells and 2 × 10^5^ sorted CD11b^+^Ly-6C^+^ macrophages from the spleens of diabetic mice or those treated with TSA and diabetes monitored (*P*=0.003). Some mice received only macrophages isolated from diabetic mice or TSA-treated mice. Four mice per group were tested. Representative data from two independent experiments are shown. Statistical significance between specified groups was assessed using a two-tailed, unpaired *t*-test. (**b**) Wild-type female NOD mice were injected with purified 2 × 10^5^ Ly6-C^+^ macrophages from diabetic mice or those treated with TSA at 13 and 21-wks of age, as indicated by the arrows. Diabetes was monitored weekly. Five mice per group were tested. Data shown are from two independent experiments.

## 4. Discussion

Although many intervention strategies have been previously reported to delay the diabetes onset in female NOD mice, the success of translating these studies into T1D treatment remains extremely slim [28]. We have reported that TSA treatment just prior to the onset of diabetes provided irreversible protection against diabetes in female NOD mice [10–13]. Protection was associated with increased histone H3 hyperacetylation in both the spleen and pancreas [10]. We now show that diabetes protection is associated with reduced infiltration of CD4^+^, CD8^+^, and Ly6-C^+^ cells in the pancreas (Fig. 1), indicating specific action of the chromatin modifier on specific immune cell types. This is consistent with reduced transcription and expression of IFN-γ and IL-17A in the peripheral lymphoid organ following TSA treatment.* These data are congruent with the notion that the protective effects of the histone modifier against autoimmune diabetes involves reduction in the generation of T helper subsets and compromised homing of the macrophages and T-cell subsets to the islets of Langerhans.

For the first time, we show herein that diabetogenesis involves exaggerated expression of *Hdac4*, *Hdac8*, and *Hdac9* genes in the secondary lymphoid organ, which were repressed by the histone modifier (Fig. 5a). Although many *Hdac* genes were also exaggerated in the pancreas of diabetic mice, surprisingly, they were resistant to TSA treatment (Fig. 5b). This was not due to the lack of access of TSA to the pancreas since *Hdac4* was enhanced in the pancreas of drug treated mice. Thus, the *Hdac* genes are differentially regulated by histone hyperacetylation in the lymphoid tissue *vs.* the target organ, consistent with the active role of T-cells in diabetes induction. Interestingly, we noted that the *Hdac3* gene was not repressed either in the spleen or pancreas by extremely low doses of TSA (Fig. 5a-b), which is at odds with the finding that administering substantial amounts of the HDAC3 selective inhibitor BRD3308 conferred marginal protection against diabetes in NOD mice [29]. It is unclear whether the BRD3308 inhibitor can exert immunosuppressive effects as TSA [10–16, Fig. 1-6]. However, BRD3308 reduced cytokine-induced apoptosis of dispersed primary islets *in vitro* and improved glycemia and insulin secretion in obese diabetic rats [30–32]. However, evidence for suppressing autoimmune responses by BRD3308 is lacking. Interestingly, TSA blocked the inflammatory cytokine-mediated beta-cell toxicity *in vitro* more effectively than the BRD3308 inhibitor [32]. Since the most potent HDAC inhibitor, TSA is endowed with both immunomodulatory [10–16, Fig. 1-6] and beta-cell protective effects [32], it seems to be an ideal drug of choice for treating autoimmune diabetes with an extremely low dose regimen compared to a large amount of the HDAC3 selective BRD3308 inhibitor required to exert marginal protection [29].

An important finding is that out of 32 immune response-related genes investigated, only a few were exaggerated during diabetes and regulated by TSA treatment. Notably, *Ifng* and *Il17a*, respectively the signature genes of Th1 and Th17 subsets, were magnified during diabetes (Fig. 2a). Although IFN-γ-producing Th1 cells were shown to mediate diabetes in neonatal NOD mice [33], the genetic absence of IFN-γ delayed but did not prevent diabetes [34]. Paradoxically, a protective role of IFN-γ in diabetes was also reported [35–36]. Thus, the role of IFN-γ in diabetes remains controversial. Whereas IL-17A was shown to contribute to diabetes [37], others raised questions about the independence of Th17 cells to induce diabetes [38–39]. Interestingly, T1D onset was shown to depend on both IL-17 and IFN-γ signaling [40]. Our data showing the repression of both *Il17a* and *Ifng* in TSA-treated mice that attained normoglycemia supports this contention. A closer analysis unraveled that the TSA treatment prevented the upregulation of *Il23*, required for the generation of Th17 cells [22] (Fig. 2) and *Ccl2* implicated in diabetes [24] (Fig. 3). A novel finding is that the gene encoding the transcription factor *Ahr* [25] was accentuated in the spleen during diabetes and repressed by the histone modifier (Fig. 4), indicating a crucial for this transcription factor in autoimmune diabetes manifestation. In addition to these upregulated genes, the steady-state levels of several genes implicated in T1D such as *Il4* [41], *Il18* [42], and *Il27p28* [43] were also diminished in TSA-treated spleen (Fig. 2). Similarly, the basal level expression of the transcription factor *Rorgt*, crucial for IL-17A transcription [26], was suppressed in the spleen of drug-treated mice. Interestingly, *Il22*, *Il23*, and *Cd39* implicated in diabetes [44] were similarly repressed in the pancreas of mice cured of diabetes. Interestingly, *Foxp3*, crucial for immunoregulation [45], was decreased in the pancreas after TSA treatment (Fig. 4), indicating that the regulation of T1D is independent of CD4^+^FoxP3^+^ cells, as we reported earlier [11]. This is similar to a model of T1D delay mediated by the administration of the migration inhibitory factor antagonist that repressed Foxp3 expression [46]. Overall, our data provide supporting evidence for the contribution of previously known genes such as *Ifng*, *Il17a*, and *Ccl2*, and importantly implicate novel roles of *Il23* and *Ahr* in diabetes manifestation, more so in lymphoid tissue than in the pancreas. Some of these genes upregulated during diabetogenesis may be the effect rather than the cause of diabetes. Further work is necessary to parse the causative role of these genes in autoimmune diabetes.

The diminished diabetogenic potential of whole splenocytes [10] and the enriched T lymphocytes (Fig. 6a) by epigenetic reprogramming reiterates the importance of T-cells in diabetes*. In addition to T lymphocytes, macrophages had been proposed to play a crucial role during the effector phase of T1D [18–19]. Both antigen presentation to cytotoxic CD8^+^ T cells [18] and direct cytotoxicity on dispersed islet cells *in vitro* [19] have been attributed to macrophages. However, direct evidence for the actual participation of macrophages in diabetogenesis *in vivo* has been difficult to establish. We observed that the CD11b^+^Ly-6C^+^ macrophages from diabetic mice failed to transfer diabetes to immunodeficient mice, indicating that the macrophages do not serve as effectors of T1D (Fig. 6a), unlike proposed previously [19]. Unexpectedly, we found that TSA treatment endowed the CD11b^+^Ly-6C^+^ macrophages with the ability to suppress disease induction upon co-transfer with diabetogenic T cells into NOD.*scid* recipients (Fig. 6a). However, the transfer of Ly-6C^+^ macrophages derived from TSA-treated mice failed to suppress ongoing diabetes in wild-type NOD mice (Fig. 6b). This paradoxical finding appears similar to the failure of polarized Th2 [41] and Th17 cells [38] derived from the BDC2.5 TCR transgenic mice to transfer diabetes into neonatal wild-type NOD mice while readily transferring the disease to immunodeficient NOD.*scid* mice. These intriguing findings were attributed to the lack of endogenous immunoregulatory mechanisms in immunodeficient mice facilitating the action of transferred cells without opposition, unlike the wild-type NOD mice. Our data are also different from diabetes suppression in NOD mice mediated by the injection of M2-type macrophages polarized *in vitro* from bone-marrow-derived monocytes using a cocktail of IL-4/IL-10/TGF-β [27]. Apparently, TSA treatment did not convert macrophages into M2-type macrophages. Therefore, this strategy, transfer of macrophages from TSA-treated mice may not be useful for T1D treatment. Although histone hyperacetylation has been proposed to impact monocytes and macrophages central to tissue homeostasis and immune responses [47–48], little evidence exists about the functional modulation of macrophages under diabetic conditions. Previously, we observed that NOD mice treated with TSA and cured of T1D displayed a lower level of the pro-inflammatory gene *Cela3b* (elastase 3) [11], which was validated herein (Fig. 3a). Moreover, TSA treatment downregulated macrophage-specific genes such as *Csf2*, *Mmp12*, and *Ym1*. Further studies are required to determine whether these changes can contribute to the suppressive function of macrophages observed in the lymphopenic environment, NOD.*scid* mouse.

## Conclusions

The data presented herein and elsewhere [10–13*] indicate that the histone modifier mediated protection against T1D involves the modified expression of a select set of immune response-related genes such as *Il17a*, *Ifng*, *Ccl2*, and *Ahr* in the spleen and altered functions of T cells and macrophages. However, changes in the expression of substantial numbers of immune response genes analyzed in the pancreas of TSA-treated mice remained minimal compared to the lymphoid tissue. This is consistent with a more active role of lymphoid cells in the induction and manifestation of autoimmune diabetes. Although we have analyzed 32 immune response-related genes in the pancreas of diabetic mice and their manipulation by TSA treatment, genes pertinent to the exocrine and endocrine tissues remain to be scrutinized. Since T1D induction is dependent on T lymphocytes, our epigenetic analysis has focused on the Th1 and Th17 subsets, allowing the parsing of the roles of, respectively, *Ifng* and *Il17a*.* The disease protection afforded by TSA treatment is inversely correlated with the levels of a small number of genes mainly in the lymphoid tissue, suggesting that they could serve as biomarkers of T1D and targets for manipulation of the disease. We believe that similar analysis of multiple immune response-related genes and inflammatory genes in T1D patients may provide significant insights into disease pathogenesis.

## Abbreviations

HDAC: histone deacetylase
qRT-PCR: quantitative reverse transcriptase-mediated polymerase chain reaction
T1D: type 1 diabetes
Th: T helper
TSA: Trichostatin A

## Acknowledgements

Mark Holterman is acknowledged for the support of this work. The expert help of the Imaging Core and Flow Cytometry Facility at the University of Illinois at Chicago in acquiring confocal images and high-speed sorting of macrophages, respectively is acknowledged.

## Author contributions

CRediT roles: Sundararajan Jayaraman: Conceptualization, data curation, formal analysis, funding acquisition, methodology, project administration, resources, supervision, validation, writing original draft, review & editing.

Maria Arianas: Methodology, review & editing.

Arathi Jayaraman: Methodology, review & editing.

## Funding

This research did not receive any specific grant from funding agencies in the public, commercial, or not-for-profit sectors.

## Footnotes

* V. Patel, A. Jayaraman, S. Jayaraman, Epigenetic reprogramming ameliorates type 1 diabetes by decreasing the generation of Th1 and Th17 subsets and restoring self-tolerance in CD4^+^ T cells, under consideration of publication. 2021.

## Notes

### Competing Interest Statement

The authors have declared no competing interest.

### Summary of Updates

Minor revisions have been made to improve clarity.

