## Supplementary figures for "Epigenetic modulation of selected immune response genes and altered functions of T lymphocytes and macrophages collectively contribute to autoimmune diabetes protection" (from another version of this preprint: version 1, 2021-07-16)

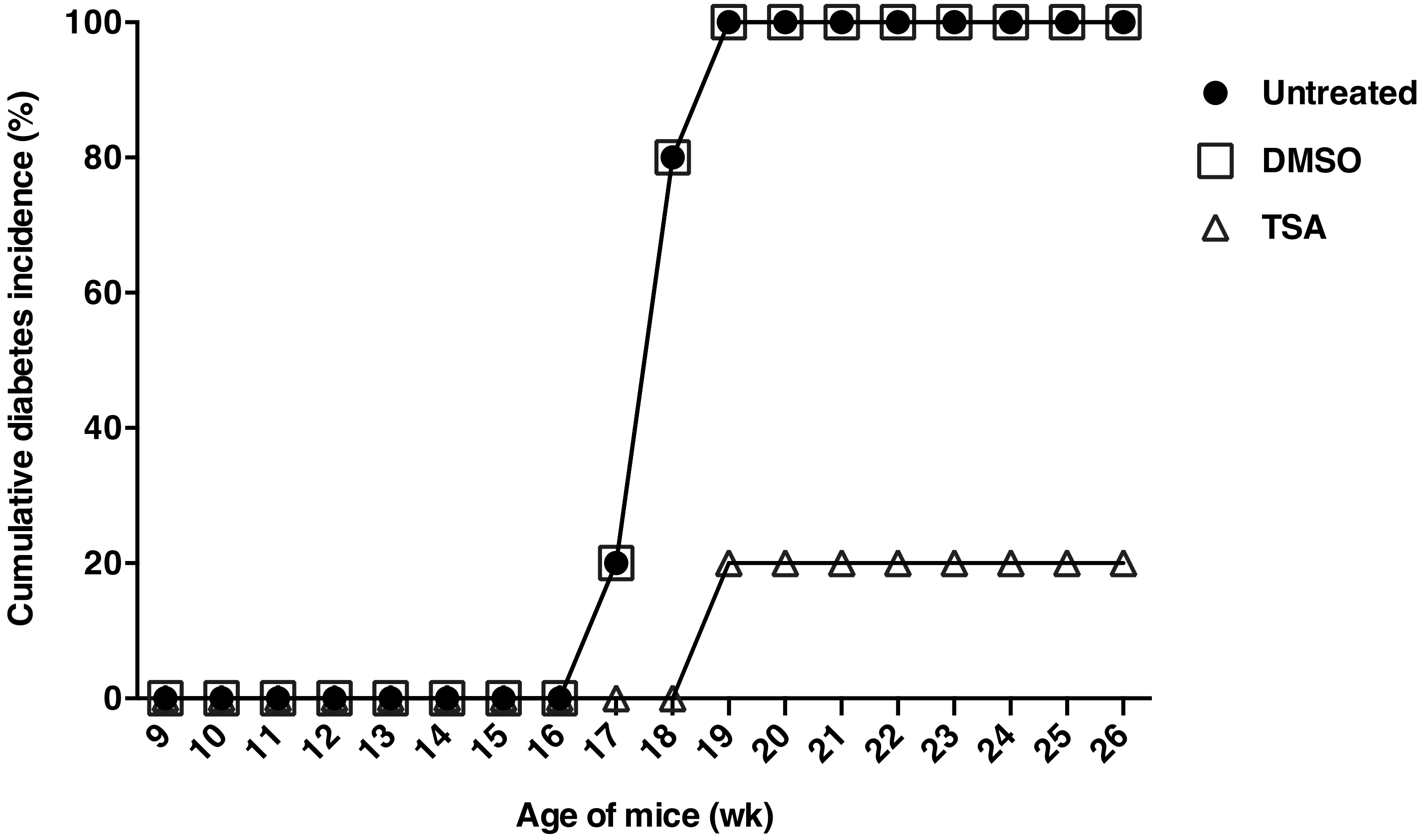


**Supplementary Fig. 1 Abrogation of T1D by TSA treatment.** Female NOD mice were treated with TSA between 16 and 24-wks of age. Controls received DMSO or left untreated. Blood glucose levels were monitored weekly, and >250 mg/dL at two consecutive weeks were considered diabetic. Five mice per group were tested in three different experiments, and the cumulative data are shown. The difference between controls (untreated and DMSO-treated mice) and those treated with TSA was statistically significant, $P<0.05$, as assessed by Wilcoxon Signed Rank Test.

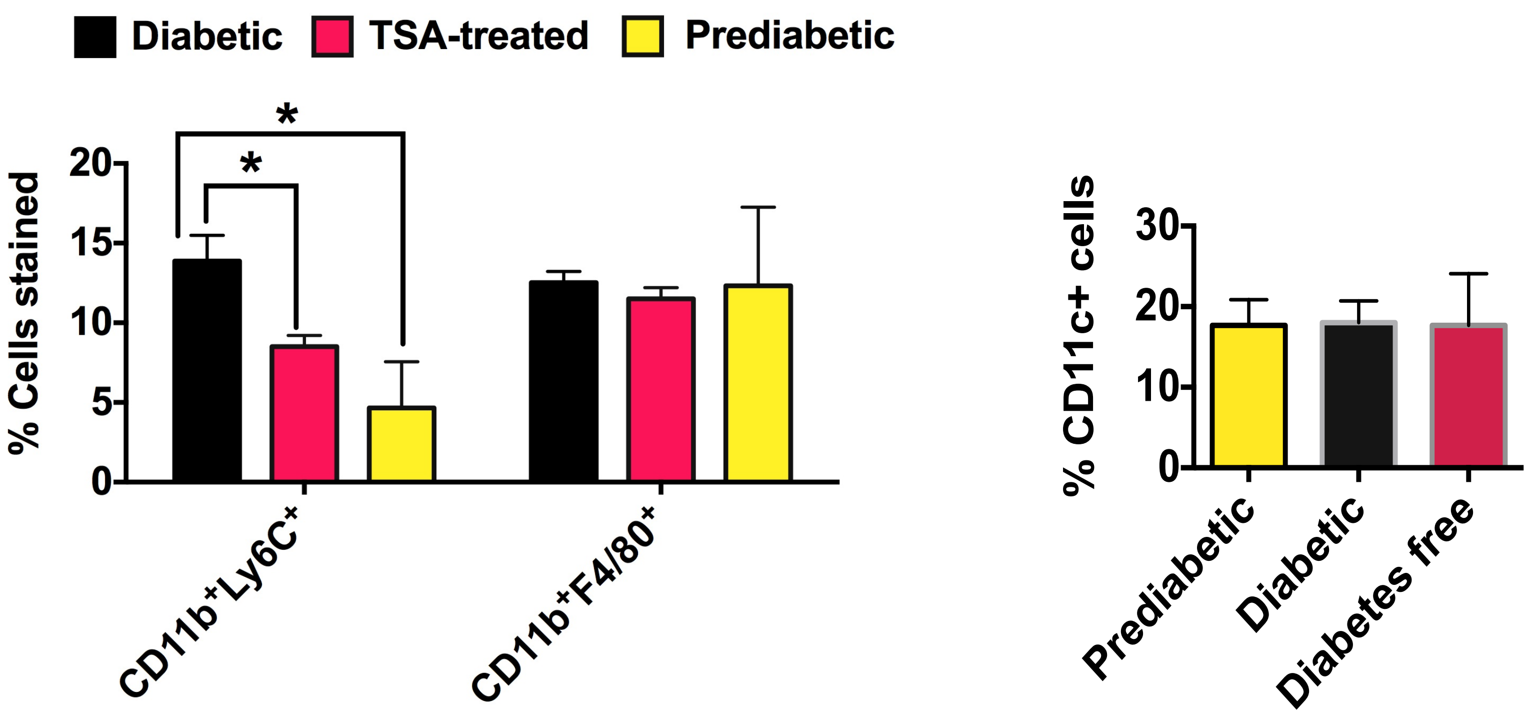


**Supplementary Fig. 2 Determination of splenic accessory cell populations by flow cytometry.** Diabetic NOD mice and those treated with TSA between 16 and 24 weeks of age were analyzed during 24-28 wks of age. Splenocytes were stained with fluoresceinated anti-CD11b, anti-Ly6C and anti-F4/80 antibodies. Cells were gated on CD11b+ cells and analyzed for the expression of Ly6C or F4/80. Viable splenocyte were analyzed for the expression of CD11c. n=5-10 per group; **P*=<0.05.
